# HdaB: a novel and conserved DnaA-related protein that targets the RIDA process to stimulate replication initiation

**DOI:** 10.1101/712307

**Authors:** Antonio Frandi, Justine Collier

## Abstract

Exquisite control of the DnaA initiator is critical to ensure that bacteria initiate chromosome replication in a cell cycle-coordinated manner. In many bacteria, the DnaA-related and replisome-associated Hda/HdaA protein interacts with DnaA to trigger the regulatory inactivation of DnaA (RIDA) and prevent over-initiation events. In the *C. crescentus Alphaproteobacterium*, the RIDA process also targets DnaA for its rapid proteolysis by Lon. The impact of the RIDA process on adaptation of bacteria to changing environments remains unexplored. Here, we identify a novel and conserved DnaA-related protein, named HdaB, and show that homologs from three different *Alphaproteobacteria* can inhibit the RIDA process, leading to over-initiation and cell death when expressed in actively growing *C. crescentus* cells. We further show that HdaB interacts with HdaA *in vivo*, most likely titrating HdaA away from DnaA. Strikingly, we find that HdaB accumulates mainly during stationary phase and that it shortens the lag phase upon exit from stationary phase. Altogether, these findings suggest that expression of *hdaB* during stationary phase prepares cells to restart the replication of their chromosome as soon as conditions improve, a situation often met by free-living or facultative intracellular *Alphaproteobacteria*.

## INTRODUCTION

All cells must be able to integrate environmental and internal cues to decide when to start the duplication of their genome. The key players, called ORC in eukaryotic and archaeal cells, and DnaA in bacteria, are conserved AAA (ATPases Associated with diverse cellular activities) proteins. In bacteria, DnaA typically binds to several so-called DnaA boxes located on a chromosomal origin and promotes the opening of the DNA double helix at a nearby AT-rich region, initiating the loading of the DNA polymerase onto the DNA ^1,2^. The correct timing of chromosome replication is directly influenced by the availability and the activity of DnaA, even in bacteria that also have additional mechanisms regulating this process ^3–5^. In most bacteria, DnaA must be bound to ATP to initiate replication and the ATPase activity of DnaA modulates its capacity to act as an initiator ^1^. In many bacteria, including the *Escherichia coli* pathogen and the environmental *Caulobacter crescentus* bacterium, the ATPase activity of DnaA is stimulated by the replisome-associated Hda/HdaA protein once the DNA polymerase is loaded onto the DNA ^6–10^. This Regulated Inactivation of DnaA (RIDA) process ensures that cells start the replication of their chromosome once and only once per cell cycle. In *C. crescentus*, the Lon ATP-dependent protease degrades DnaA-ADP more efficiently than DnaA-ATP ^11–13^, so that the RIDA process not only inactivates DnaA but also destabilizes it, providing a robust control system ^4^.

When bacteria face adverse growth conditions, they must adjust their cell cycle accordingly to ensure survival. These adaptation mechanisms are still poorly understood, although most bacteria are believed to inhibit the replication of their chromosome under such conditions. In *C. crescentus*, translation of the *dnaA* transcript is strongly inhibited in response to nutrient limitations, leading to a rapid clearance of DnaA by Lon ^14,15^. Thus, *C. crescentus* cells that enter stationary phase are mostly arrested at the pre-divisional stage of their cycle ^16^. How such cells can then exit from stationary phase, re-start the replication of their genome and proliferate again when conditions get better remains unclear, although this must be a frequent need in natural environments. In this study, we uncovered the existence of a novel and conserved DnaA-related protein that is specifically expressed during stationary phase, most likely preparing cells to initiate DNA replication during exit from stationary phase.

## MATERIALS AND METHODS

### Plasmids and strains

Oligonucleotides and plasmids used in this study are described in Table S1 and Table S2, respectively. Bacterial strains used in this study are described in Table S3. Material and methods used to construct novel plasmids and strains are described in Supplementary Information.

### Growth conditions and synchronization

*E. coli* strains were cultivated at 37°C in LB medium or on LB + 1.5% agar (LBA). *C. crescentus* was cultivated at 30°C in peptone yeast extract (PYE) complex medium or on PYE + 1.5% agar (PYEA). When required for selections or to maintain plasmids or genetic constructs, antibiotics were added at the following concentrations for solid/liquid media (μg/ml), respectively: tetracycline (Tet; PYE: 2/1, LB: 10/10), kanamycin (Km; PYE: 25/5, LB: 50/25), gentamycin (Gent; PYE: 5/1, LB: 10/10), spectinomycin (Spec; PYE: 100/25, LB: 50/50), streptomycin (Strep; PYE: 5/5, LB: 30/30) and nalidixic acid (Nal; PYEA: 20). When mentioned, glucose and/or xylose were added at a final concentration of 0.2% and/or 0.3%, respectively. When indicated, vanillate was added at a final concentration of 1mM. Synchronized cultures of *C. crescentus* were obtained by centrifugation in a Percoll (Sigma, DE) density gradient followed by isolation of swarmer cells using a protocol adapted from ^38^. Swarmer cells were then released into PYE medium for cell cycle studies.

### Immunoblot analysis

Proteins were resolved on 10% (for DnaA) or 12.5% (for HdaA, M2-tagged HdaB or CcrM) SDS-PAGE gels. Proteins were electro-transferred to PVDF membranes (Merck Millipore, MA, USA). Immuno-detection was performed with polyclonal antibodies. Anti-DnaA ^18^, anti-HdaA ^8^, anti-CcrM ^39^ and anti-rabbit conjugated to horse-radish peroxidase (HRP) (Sigma, GER) sera were diluted 1:10000. Anti-GFP (Thermo Fischer Scientific, USA) and anti-FLAG (Sigma, GER) sera were diluted 1:10000. Anti-mouse conjugated to HRP (Promega, USA) was diluted 1:5000. Immobilon Western Blotting Chemo-luminescence HRP substrate (Merck Millipore, MA, USA) was used and membranes were scanned with a Fusion FX (Vilber Lourmat, FR) instrument to detect the chemiluminescent signal. Relative band intensities on images were quantified using the ImageJ software.

### β-galactosidase assays

β-galactosidase assays were done using a standard procedure as previously described ^40^.

### Microscopy and image analysis

Cells were immobilized on a slide using a thin layer of 1% agarose dissolved in dH_2_O and imaged immediately. Phase contrast (Ph3) and fluorescence images were taken using the microscope system described in ^41^. The MicrobeJ software ^42^ was then used: (i) to estimate the proportion of dead cells on fluorescence images of LIVE/DEAD-stained cells using default parameters of the Bacteria mode of detection option; (ii) to measure the length of cells on phase-contrast images using the Medial-Axis mode of detection option; (iii) to detect and count fluorescent foci in cells on fluorescence images using the Foci-mode of detection option (foci were counted if their signal intensity was minimum two fold above the signal of the cell cytoplasm).

### qPCR analysis to evaluate *Cori/ter* ratios

Chromosomal DNA was extracted from cells using the PureGene genomic DNA extraction kit (QIAGEN, GER). qPCR reactions were performed in a final volume of 10 μl containing: 1 μl of genomic DNA at 0.1 ng/μl, 5 μl of 2× GoTaq q-PCR master mix (Promega, USA) and 0.6 μM of AF77/78 or AF79/80 primer pairs targeting a region close to the *Cori* or to the terminus (*ter*), respectively. qPCR cycles were run as described elsewhere ^6^ but using a Rotor Gene-Q instrument (QIAGEN, GER). To quantify the relative abundance of *Cori* and *ter* chromosomal regions in each sample, a calibrator-normalized relative analysis was performed.

### Co-immunoprecipitation assays

Cells were harvested by centrifugation at 6000×*g* for 20 min. at 4 °C, washed with 25 ml of PBS (Phosphate Buffered Saline pH 7.4) and centrifuged again at 6000×*g* for 10 min. at 4 °C. Cells were then resuspended in 10 ml of PBS supplemented with 2 mM DTSP (3,3′-Dithiodipropionic acid di(N-hydroxysuccinimide ester dissolved in DMSO, a crosslinking agent from Sigma, MO, USA) and incubated for 30 min. at 30 °C with shaking. To stop the crosslinking reaction, 1.5 ml of 1M Tris-HCl (150 mM final) was added and incubation at 30°C with shaking was continued for 30 more min. Cells were then centrifuged at 6000×*g* for 10 min at 4°C, washed with 30 ml PBS and centrifuged at 6000×*g* for 10 min at 4°C. The resulting pellet was carefully resuspended in 20 ml CoIP buffer (20mM Hepes, 150mM NaCl, 20% Glycerol, 80mM EDTA pH 8 and complete EDTA-free protease inhibitor cocktail according to instructions (Roche, CH)), centrifuged at 6000×*g* for 20 min. at 4°C and resuspended in 2 ml of NP40 lysis buffer (1× CellLytic B (Sigma, MO, USA), 50 U DNAse I (Fermentas, MA, USA), 75000 U ReadyLyse (Epicentre, WI, USA), 10 mM MgCl_2_, 2% NP40 (Merck Millipore, MA, USA), 1× protease inhibitor cocktail) and incubated at 25°C for 30 min. with end-over-end rotation. Cells were then subjected to five cycles of sonication on ice (10 sec. pulse/1 min. pause on ice) and centrifuged at 13362×*g* for 15 min. at 4°C. The resulting cell extract (INPUT) was used to determine the total protein concentration using Bradford assay (Biorad, USA). 10 mg of total proteins were mixed with Magnetic anti-FLAG beads (Sigma, USA) and incubated over-night with end-over-end rotation at 4°C. A small aliquot of the supernatant (SN) was collected at that step. Suspensions were then washed 6 times with 1mL of CoIP buffer + 1% Triton X100. Lastly, beads were resuspended in 1× Laemmli Sample Buffer (Biorad, USA) and incubated at 70°C for 10 min (IP). Samples (INPUT/SN/IP) were heated at 95°C for 10 min. in 1× Læmmli Sample Buffer for immunoblotting.

### Genetic screen to isolate suppressors of the toxicity associated with *hdaB* expression in *C. crescentus*

pHPV414 is a *himar1* transposon (*Tn::km*) delivery plasmid ^43^ and it was introduced by conjugation into NA1000 cells carrying the pBX-HdaB^Ae^ plasmid and cultivated into PYE + glucose medium. More than 8’000 random mutants were then picked and transferred in parallel onto PYEA plates containing either glucose or xylose. Only two transposon mutants were found to grow on both media; these were named JC 2077 and JC2078. For each of these two mutants, the pBX-HdaB^Ae^ plasmid was extracted and introduced back into NA1000 cells, to confirm that presence of these plasmids still led to toxicity in *WT* cells, thereby indicating that chromosomal mutations in JC2077 and JC2078 suppress the toxicity associated with *hdaB*^*Ae*^ expression from pBX-HdaBAe. Transposon mutations from strains JC2077 and JC2078 were transduced into NA1000 cells using ФCr30 and kanamycin selection, giving JC2080 and JC2079, respectively. Strains JC2080 and JC2079 were then used to map the transposon insertion sites in the chromosome of these mutants of interest. Their chromosomal DNA was extracted and digested with SacII. DNA fragments were then self-ligated before transformation of *E. coli* S17-1 *λpir* with the ligation mix. The transposon insertion sites were then identified by sequencing using primer himup2 ^43^ and comparison with the NA1000 genome available on NCBI (NC_011916): strain JC2080 carried a *dnaK::Tn* insertion at position 9955 of this genome, while strain JC2079 carried a *recQ::Tn* insertion at position 3735008 of this genome. To further verify that *dnaK::Tn* or *recQ::Tn* insertions were sufficient to suppress the toxicity associated with the expression of different HdaB proteins, pBX-HdaB plasmids were introduced back into JC2079 and JC2080 prior to phenotype analysis (Fig. 2, 3).

## RESULTS

### HdaB is a conserved DnaA-related protein

We noticed that the majority of *Alphaproteobacteria* that possess a HdaA protein also encode a second protein that shows similarities with DnaA (Fig. 1a) and thus logically named such proteins HdaB (homologous to DnaA B). HdaB proteins are short proteins resembling the domain IV of DnaA proteins. One example is the uncharacterized CCNA_01111 protein of the *C. crescentus* NA1000 strain (HdaB^Cc^), which displays 32% of identity with DnaA^Cc^ domain IV (Fig. 1b and Supplementary Fig. 1). Considering the DnaA domain IV is involved in DNA binding and in interactions with Hda/HdaA during the RIDA process ^2,17^, we suspected that HdaB proteins might be involved in the control of chromosome replication in certain *Alphaproteobacteria*. Indeed, we now provide evidences that HdaB proteins from minimum three different *Alphaproteobacteria* can interfere with the RIDA process and affect replication frequency in the *C. crescentus* model bacterium.

**Figure 1:**
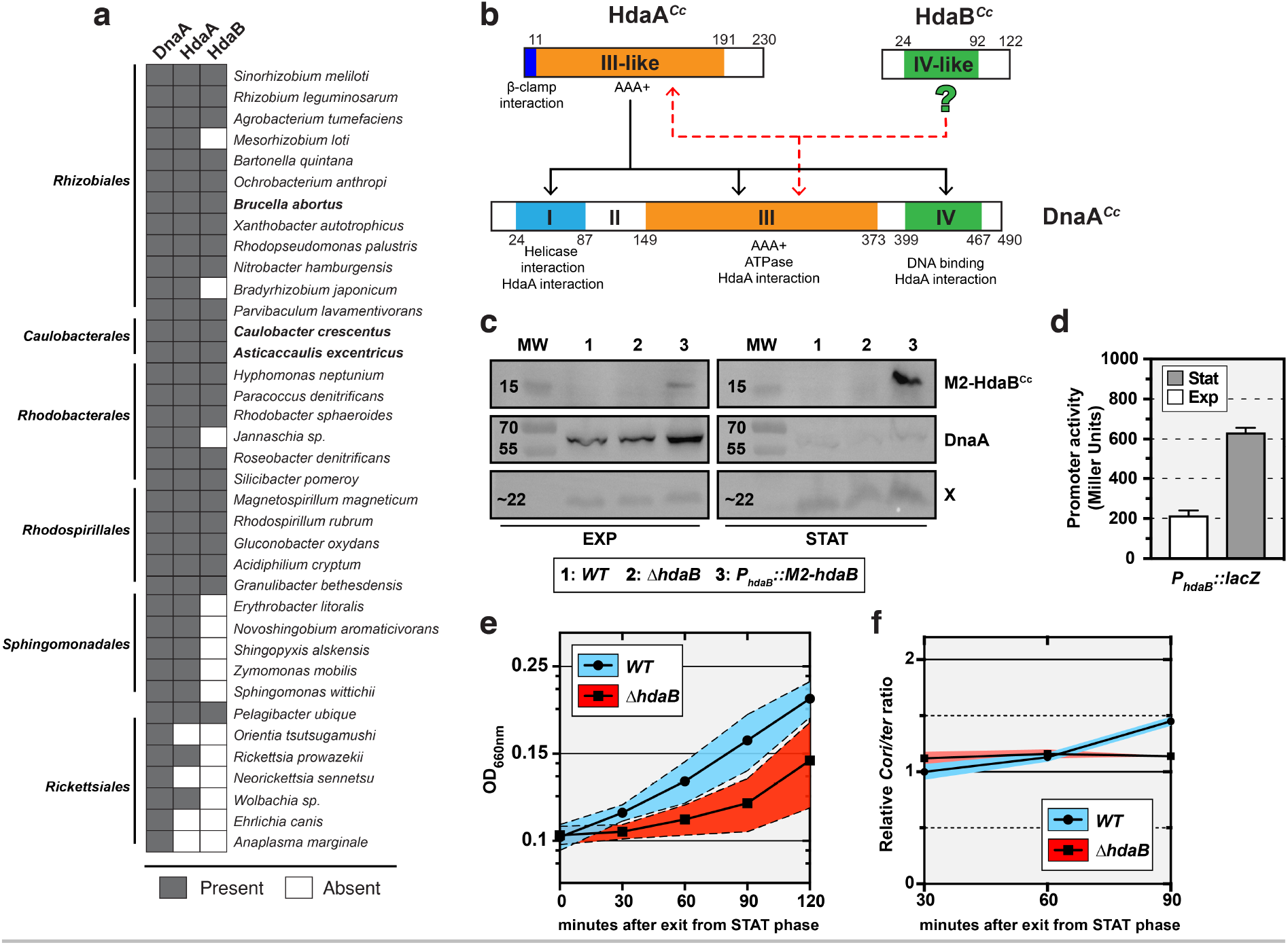
HdaB is a conserved DnaA-related protein that accumulates in stationary phase. **(a)** Distribution of DnaA, HdaA and HdaB proteins in *Alphaproteobacteria*. The presence or the absence of a homolog of each protein was estimated using bi-directional best-blast-hit (BBH) using the *C. crescentus* genome as reference. HdaB from species in bold have been used in this study. **(b)** Similarities between DnaA^Cc^ domains and HdaA^Cc^ or HdaB^Cc^. The predicted (from ^2^) or demonstrated (from ^6–9^) function of each protein motif or domain is indicated under its position. Black arrows indicate interactions demonstrated between Hda^Ec^ and DnaA^Ec^ in *E. coli* (^2^). Red dashed arrows indicate putative interactions between HdaB^Cc^ and HdaA^Cc^ or DnaA^Cc^. Numbers indicate amino-acid positions on each protein. **(c)** Immunoblots showing the intracellular levels of M2-HdaB^Cc^ and DnaA^Cc^ in *C. crescentus* cells cultivated in exponential (EXP) or stationary (STAT) phase. Strains NA1000 (*WT*), JC1456 (Δ*hdaB*) and JC2106 (*P_hdaB_::M2-hdaB*) were cultivated in PYE medium and samples were collected at an OD_660nm_ of ∼0.3 (EXP) and after an over-night culture (STAT). Proteins were analyzed by SDS-PAGE and detected by immunoblotting using anti-M2 (for M2-HdaB) or anti-DnaA (for DnaA and the unknown X protein that serves as a loading control here) antibodies. MW stands for the approximate molecular weight in kD. **(d)** Activity of the *hdaB*^*Cc*^ promoter in cells cultivated in exponential or stationary phase. The p*lacZ290*-P*hdaB* plasmid was introduced into the NA1000 (*WT*) strain. Cells were then cultivated in PYE medium and β-galactosidase assays were performed when cultures reached an OD_660nm_ of ∼0.3 (EXP) and after an over-night incubation (STAT). Error bars correspond to standard deviations from minimum three experiments. **(e)** Growth of a Δ*hdaB* mutant upon exit from stationary phase. *C. crescentus* strains NA1000 and JC1456 (Δ*hdaB*) were cultivated into PYE medium until stationary phase (over-night) and cultures were diluted back into fresh PYE medium to reach an OD660nm of 0.1 at time 0. Blue or red shaded areas represent standard deviations from 4 independent cultures of each strain. **(f)** *Cori/ter* ratio in Δ*hdaB* cells during exit from stationary phase. *C. crescentus* strains NA1000 and JC1456 (Δ*hdaB*) were cultivated into PYE medium until stationary phase (over-night) and cultures were diluted back into fresh PYE medium to reach an OD660nm of 0.1 at time 0. gDNA was extracted at the indicated time points and *Cori/ter* ratio were estimated by qPCR. Results were then normalized so that the *Cori/ter* ratio of *WT* cells at time 30 min. equals 1. Blue or red shaded areas represent standard deviations from two independent cultures of each strain.

### HdaB accumulates in stationary phase *C. crescentus* cells

To get a first indication on the potential role of HdaB proteins during the life cycles of *Alphaproteobacteria*, we compared the expression of HdaB^Cc^ in *C. crescentus* cells cultivated in different growth conditions. We used cells expressing HdaB^Cc^ with a M2 (FLAG) epitope at its N-terminus (M2-HdaB^Cc^) from the native *hdaB* promoter to facilitate detection by immunoblotting. Surprisingly, we found that M2-HdaB^Cc^ levels were hardly detectable in exponentially-growing cells, while accumulating at much higher levels in stationary-phase cells (Fig. 1c). This observation suggested that the synthesis of HdaB^Cc^ may be stimulated upon entry into stationary phase. To test this possibility more directly, we constructed a plasmid carrying a reporter where *lacZ* transcription is under the control of the *hdaB*^*Cc*^ promoter region and introduced this plasmid into *C. crescentus* cells. Cells were then cultivated from exponential to stationary phase and β-galactosidase assays were performed to compare the activity of the *hdaB*^*Cc*^ promoter. Consistent with our previous findings (Fig. 1c), we found that the *hdaB*^*Cc*^ promoter was ∼three-fold more active in stationary phase cells than in exponentially growing cells (Fig. 1d). Altogether, these results suggested that HdaB^Cc^ could play a role during stationary phase or during exit from stationary phase, rather than during exponential growth when HdaB levels remain extremely low. Consistent with this proposal, we found that an in-frame deletion of the *hdaB*^*Cc*^ gene did not affect the generation time of *C. crescentus* cells during exponential growth (96.8 ± 6.6 minutes for Δ*hdaB* cells, compared to 95.2 ± 10.7 minutes for wild-type cells in complex PYE medium). Instead, deletion of the *hdaB* gene significantly expanded the duration of the lag phase during exit from stationary phase (Fig. 1e). Interestingly, this growth delay appeared to coincide with a delay in replication restart since the *origin/terminus* (*Cori/ter*) ratio estimated by quantitative real-time PCR (qPCR) increased earlier in wild-type than in *ΔhdaB* populations of cells (Fig. 1f).

### Ectopic expression of HdaB proteins inhibits cell division and leads to cell death

Considering that *hdaB*^*Cc*^ is hardly expressed in fast-growing cells, we anticipated that looking at the impact of *hdaB* expression in fast-growing cells may turn out more informative than characterizing *hdaB* deletion strains. Thus, we cloned *hdaB* genes from three different *Alphaproteobacteria* under the control of the xylose-inducible *xylX* promoter (*Pxyl*) of *C. crescentus* on a medium copy number plasmid and introduced these plasmids into wild-type *C. crescentus* cells. Strikingly, in all three cases, actively growing cells expressing *hdaB* homologs became elongated with frequent failure during the division process compared to control cells (Fig. 2a and Supplementary Fig. 2a), showing that expression of *hdaB^Cc^*, *hdaB*^*Ae*^ (from *Asticaccaulis excentricus*) or *hdaB*^*Ba*^ (from *Brucella abortus*) interferes directly or indirectly with cell division. Moreover, cell elongation upon *hdaB* expression rapidly lead to cell death. In liquid medium, up to ∼ 50% of the cells died after just four hours of induction (Supplementary Fig. 2b). On solid medium containing the xylose inducer, very few cells expressing *hdaB*^*Cc*^ or *hdaB*^*Ba*^ could form visible colonies, while nearly no cell expressing *hdaB*^*Ae*^, except potential suppressors, could form visible colonies (Fig. 2b), demonstrating the strong impact that an ectopic expression of *hdaB* homologs can have on *C. crescentus* proliferation. Importantly, none of these phenotypes were observed when a randomly isolated mutant HdaB^Cc^(A58T) protein was expressed in identical conditions (Fig. 2b and Supplementary Fig. 2b,c), showing that they were not simply connected to a proteotoxic stress generated by protein over-production.

**Figure 2:**
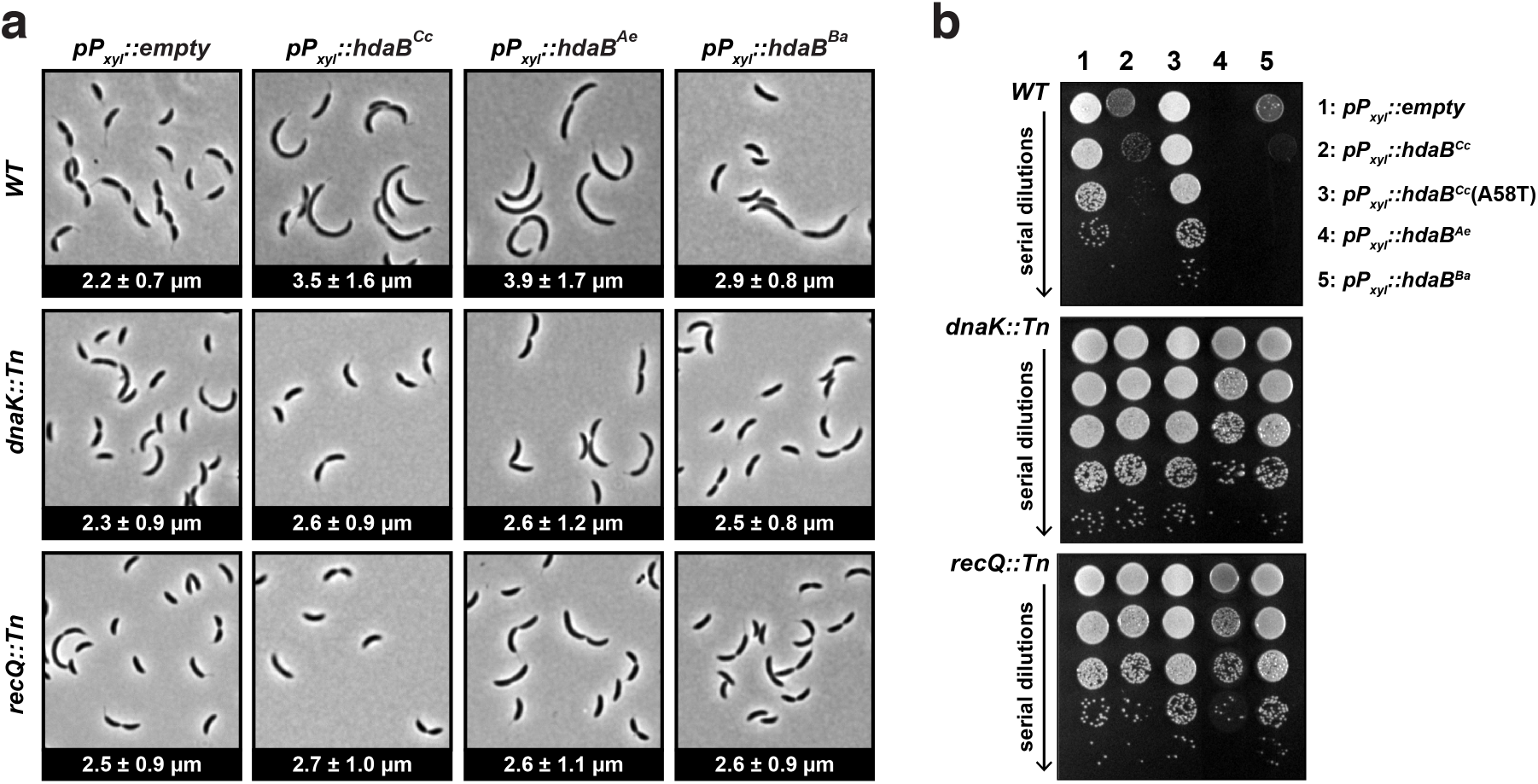
*hdaB* expression interferes with cell division and leads to cell death in *C. crescentus*. **(a)** Cellular morphology of *C. crescentus* cells expressing *hdaB*^*Cc*^ from *C. crescentus*, *hdaB*^*Ae*^ from *A. excentricus* or *hdaB*^*Ba*^ from *B. abortus*. Medium copy number vectors (pBXMCS-4 derivatives) expressing or not *hdaB* genes from a P*xyl* promoter were introduced into NA1000 (*WT*), JC2080 (*dnaK::Tn*) or JC2079 (*recQ::Tn*) cells. The resulting strains were cultivated into PYEG medium until mid-exponential phase and xylose was added for 4 hours before microscopy images were acquired. The numbers indicated at the bottom of each image correspond to the median medial axis length of cells in μm with standard deviations from three independent experiments. **(b)** Growth of colonies of *C. crescentus* cells expressing *hdaB*^*Cc*^, *hdaB*^*Cc*^(*A58T*), *hdaB*^*Ae*^ or *hdaB*^*Ba*^. The strains described in panel **(a)** were cultivated in PYE medium until mid-exponential phase and cultures were serially diluted (10-fold each time) before plating onto PYEA medium containing xylose to induce expression of *hdaB* genes.

### Expression of HdaB proteins leads to over-initiation of chromosome replication in growing cells

Since HdaB shows similarities with the domain IV of DnaA (Fig. 1b and Supplementary Fig. 1) and since its ectopic expression leads to cell division defects and cell death (Fig. 2 and Supplementary Fig. 2), like cells displaying DNA replication defects ^6,8,9,18^, we hypothesized that HdaB may play a direct role in replication control. To test this possibility, we first introduced the previously described plasmids expressing *hdaB* homologs from the P*xyl* promoter into a *C. crescentus* strain where the *parB* gene was exchanged with a *cfp-parB* construct, expressing a functional CFP-tagged ParB protein ^19^. ParB proteins bind to *parS* sequences, which are located next to the *Cori*. Thus, CFP-ParB can be used as a *proxy* to visualize *Cori* in single cells by fluorescence microscopy: in *C. crescentus*, it forms a single fluorescent focus at the flagellated cell pole before the onset of DNA replication and then forms two fluorescent foci located at each cell pole right after the onset of DNA replication ^19^ (Fig. 3b, *empty vector images*). When cells were cultivated with the xylose inducer for four hours, we observed that cells expressing each of the three *hdaB* homologs often accumulated more than two foci per cell (up to six foci per cell), which was only rarely observed with cells carrying the empty control vector or expressing the inactive HdaB^Cc^(A58T) protein (Fig. 3a). This first observation showed that the initiation of DNA replication can still take place in actively growing *C. crescentus* cells expressing *hdaB* homologs, but that the control of its frequency seems disturbed, leading to over-initiation. To bring more light on this observation, we also isolated G1 swarmer cells from the *cfp-parB* strains carrying the empty vector or the *hdaB^Cc^*-expressing plasmid and followed their differentiation into S-phase stalked cells and their division during time-lapse fluorescence microscopy experiments. Strikingly, as early as 20 minutes after the addition of the xylose inducer, when cells differentiated into stalked cells, we already observed many cells with more than two CFP-ParB foci when *hdaB*^*Cc*^ was expressed (Fig. 3b). While control cells typically displayed only two CFP-ParB foci during the S-phase of the cell cycle, cells expressing *hdaB*^*Cc*^ displayed an average of approximately four distinct CFP-ParB foci, showing that such cells can restart DNA replication multiple times during the same cell cycle. To further confirm that *hdaB*^*Cc*^ expression leads to over-initiation, we also measured the *Cori/ter* ratio by qPCR using un-synchronized populations of cells and found that it increased by a factor of ∼1.6 in cells expressing *hdaB*^*Cc*^ for four hours compared to control cells (Fig. 3c).

**Figure 3:**
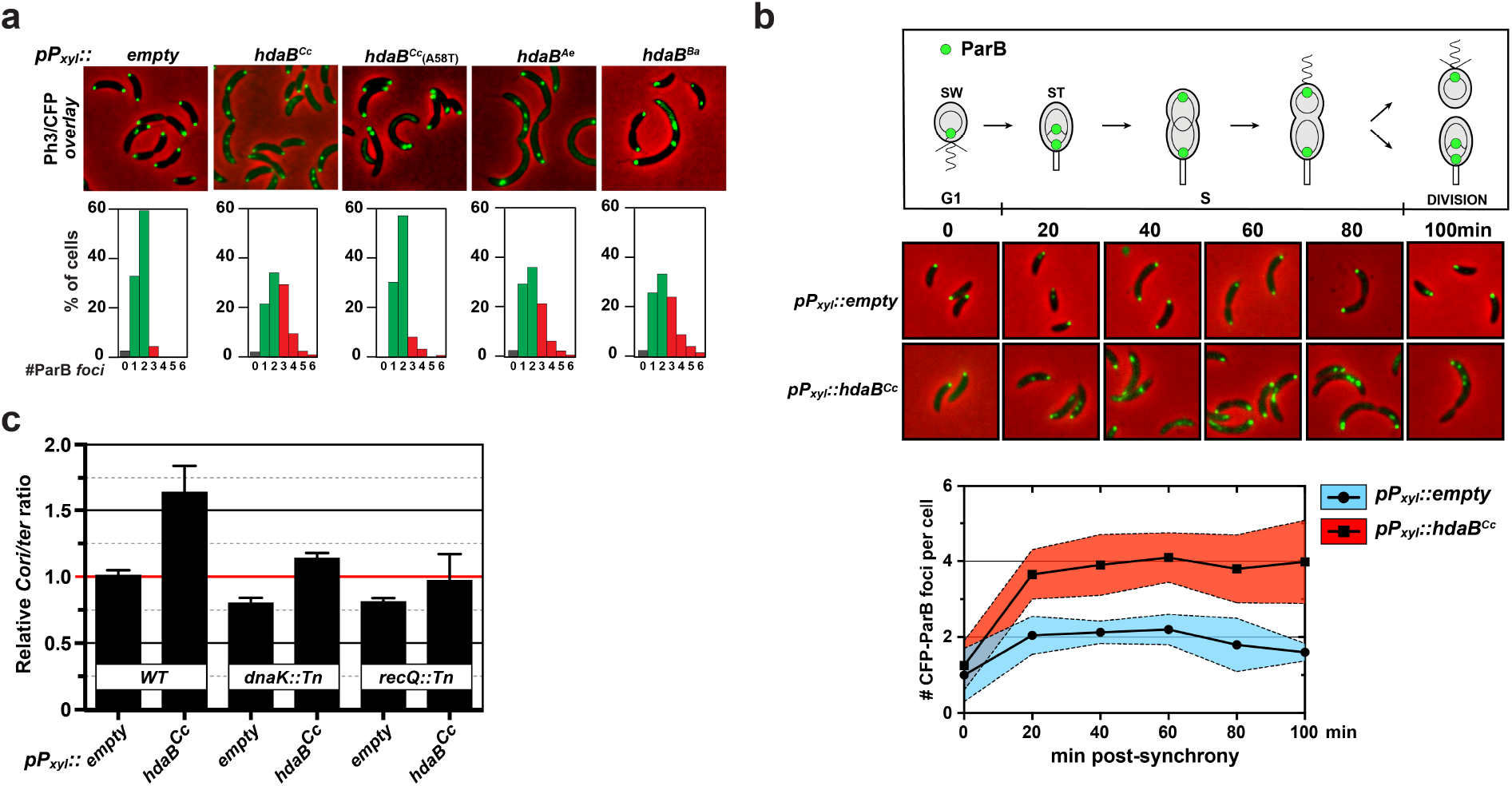
*hdaB* expression leads to fast over-replication in *C. crescentus*. **(a)** Microscopy detection of ParB-CFP foci in *C. crescentus* cells expressing *hdaB* genes. Vectors expressing or not *hdaB^Cc^, hdaB^Cc^(A58T)*, *hdaB*^*Ae*^ or *hdaB*^*Ba*^ from the *Pxyl* promoter were introduced into the MT190 strain (*P_parB_::cfp-parB*). The resulting strains were cultivated into PYEG medium to mid-exponential phase and xylose was then added for 4 more hours prior to fluorescence microscopy. Representative images corresponding to phase contrast and fluorescence (colored in green on images) overlays are shown. The number of CFP-ParB foci per cell was estimated using ∼200 cells on multiple images acquired using each strain. Results are shown as a percentage of cells displaying each number of CFP-ParB foci in each population and are averages of minimum three independent experiments. **(b)** Time-course microscopy images to detect CFP-ParB foci in isolated swarmer cells expressing *hdaB^Cc^*. MT190 strains carrying the empty vector or the vector expressing *hdaB*^*Cc*^ from *Pxyl* were cultivated until mid-exponential phase in PYEG medium before synchronization. Isolated swarmer cells were released into fresh PYE medium containing xylose and Ph3/CFP images (overlays are shown) were acquired at the indicated time points. The number of CFP-ParB foci per cell was estimated using ∼200 cells acquired using each strain at each time point. The blue or red-shaded areas correspond to standard deviations from two independent cultures of each strain post-synchrony. **(c)** Comparison of the *Cori/ter* ratio in populations of NA1000 (*WT*), JC2080 (*dnaK::Tn*) or JC2079 (*recQ::Tn*) cells expressing or not *hdaB^Cc^*. *Cori/ter* ratios were measured by qPCR using genomic DNA extracted from cells cultivated to mid-exponential phase in PYEG medium and in the presence of the xylose inducer for 4 hours. Results were normalized so that the *Cori/ter* ratio of *WT* cells carrying the empty vector equals 1 (highlighted with a red line).

Altogether, these results demonstrated that HdaB homologs from three different *Alphaproteobacteria* can disturb the frequency of the initiation of chromosome replication in *C. crescentus*, which is normally controlled by the RIDA process.

### HdaB interacts with HdaA to inhibit the RIDA process

In *C. crescentus*, the RIDA process controls DnaA activity and DnaA levels, by stimulating the ATPase activity of DnaA and its subsequent degradation by ATP-dependent proteases ^4,6,8,9,11^. It is mediated by the conserved HdaA protein ^7,8^. In *E. coli*, the RIDA process starts with an interaction between the DnaA^Ec^ domain IV and Hda, which, in turn, promotes the interaction between the AAA+ domain of the two proteins ^2,17^. Considering that HdaB proteins show similarities with the DnaA domain IV (Fig. 1b and Supplementary Fig.1), we hypothesized that HdaB could interact with HdaA to block the RIDA process in *Alphaproteobacteria*, leading to higher intracellular levels of active DnaA and subsequent over-initiation events leading to cell death. Consistent with this proposal, we observed that the Leu-422 residue of the DnaA^Ec^ domain IV, which is critical for the DnaA^Ec^/Hda interaction ^17^, is present in HdaB^Cc^ (Supplementary Fig.1), while the Arg-399 residue, which is necessary for DnaA^Ec^ binding to DnaA boxes ^20^, is not.

As a first step to test this hypothesis, we probed whether HdaA and HdaB can interact with one another in *C. crescentus* cells and whether this interaction is required for the toxic function of HdaB. To do so, we first engineered two constructs expressing either M2-HdaB^Cc^ or HdaB^Cc^-M2 from the *Pxyl* promoter into a medium copy number plasmid, introduced these constructs into wild-type *C. crescentus* cells and verified the functionality of M2-tagged HdaB^Cc^ proteins compared to HdaB^Cc^. We found that the ectopic expression of M2-HdaB^Cc^ (Fig. 4 and Supplementary Fig. 2) in the presence of xylose for four hours had as much impact on cell viability and cell morphology as that of HdaB^Cc^ (Fig. 2 and Supplementary Fig. 2), while expression of HdaB^Cc^-M2 had no impact (Fig.4 and Supplementary Fig. 2). This result showed that the addition of a M2 tag at the C-terminus of HdaB^Cc^ interferes with the function of HdaB, while a M2 tag at its N-terminus does not appear to impact its function. We then used these two strains to perform anti-M2 immunoprecipitation experiments followed by anti-HdaA immunoblots to get a first insight on the potential interaction between native HdaA and M2-tagged HdaB proteins. We found that the functional M2-HdaB^Cc^ protein co-immunoprecipitated with HdaA, while the non-functional HdaB^Cc^-M2 protein did not (Supplementary Fig. 3). Since control cells lacking *hdaA* could not be used in such experiments as *hdaA* is an essential gene ^8^, we also engineered two more constructs expressing either the functional M2-HdaB^Cc^ or the non-functional HdaB^Cc^-M2 from the vanillate-inducible *Pvan* promoter into a medium copy number plasmid and introduced these plasmids into a *C. crescentus* strain expressing a functional GFP-tagged HdaA protein from the native chromosomal P*xyl* promoter ^7,8^. These strains were then cultivated in the presence of the xylose and vanillate inducers for four hours and cell extracts were used to perform anti-M2-immunoprecipitation experiments followed by anti-GFP immunoblots. We found that the functional M2-HdaB^Cc^ protein co-immunoprecipitated with the functional GFP-HdaA protein, while the non-functional HdaBCc-M2 protein did not (Fig. 4). Worth noticing, none of the M2-tagged HdaB^Cc^ proteins co-precipitated with DnaA, suggesting that HdaB^Cc^ does not target DnaA directly. Altogether, these results provide a strong indication that HdaB must interact with HdaA to exert its toxic function.

**Figure 4:**
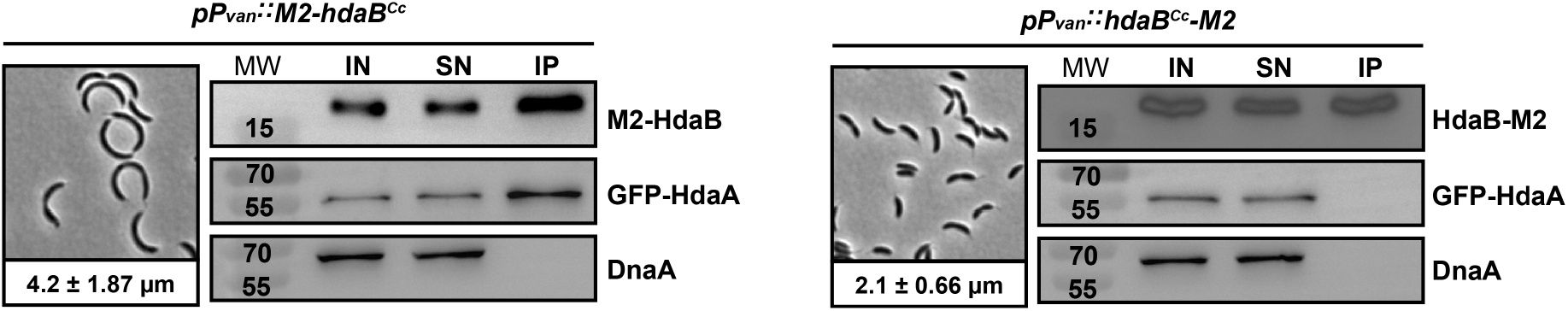
HdaB interacts with HdaA but not DnaA in *C. crescentus.* Left panels: Cellular morphology of NA1000 (*WT*) *C. crescentus* cells expressing *M2-hdaB*^*Cc*^ or *hdaB^Cc^-M2* from the *Pxyl* promoter on the pBXMCS-4 vector. The resulting strains were cultivated into PYEG medium until mid-exponential phase and xylose was added for 4 hours before microscopy images were acquired. The numbers indicated at the bottom of each image correspond to the median medial axis length of cells in μm with standard deviations from three independent experiments. Right panels: Probing the *in vivo* interaction(s) between M2-HdaB^Cc^/HdaB^Cc^-M2 and GFP-HdaA^Cc^/DnaA by co-immunoprecipitation. JC208 (*P*_*xylX*_*::gfp-hdaA*) cells carrying pBV-M2-HdaB^Cc^ or pBV-HdaB^Cc^-M2 were cultivated to mid-exponential phase in PYEG medium. Xylose and vanillate were then added into the medium for 4 hours (for *gfp-hdaA* and *M2-tagged hdaB*^*Cc*^ expression, respectively) before cell lysis and immunoprecipitation using anti-M2 antibodies. Images show immunoblots obtained using anti-GFP, anti-DnaA or anti-M2 antibodies. IN: cell extract/input; SN: non-precipitated proteins/supernatant; IP: precipitated proteins/immunoprecipitation. MW: molecular weight in kD.

As a second step to test this hypothesis, we probed whether the ectopic expression of HdaB proteins leads to higher DnaA accumulation in *C. crescentus*, as expected if they target HdaA and inhibit the RIDA process. We performed immunoblot experiments using extracts of *C. crescentus* cells expressing any of the three selected *hdaB* homologs of *Alphaproteobacteria* from a *Pxyl* promoter on a plasmid and found that DnaA accumulated at higher intracellular levels in all three cases, compared to control cells carrying the empty vector (Fig. 5a). Four hours after the addition of xylose into the medium, cells expressing *hdaB* homologs accumulated at least 2.5-fold more DnaA than control cells (Fig. 5a). Since the RIDA process inhibits DnaA accumulation by promoting its degradation, we also compared the stability of DnaA in *hdaB*-expressing cells and in control cells by immunoblotting using samples collected at different times after an addition of chloramphenicol to stop DnaA synthesis (Fig. 5b). We found that the half-life of DnaA roughly doubled (from ∼40 minutes to ∼80 minutes) when HdaB proteins were expressed from *Pxyl* for four hours, compared to control cells. Importantly, a similar expression of the inactive HdaB(A58T)^Cc^ protein had essentially no impact on DnaA stability (Fig. 5b).

**Figure 5:**
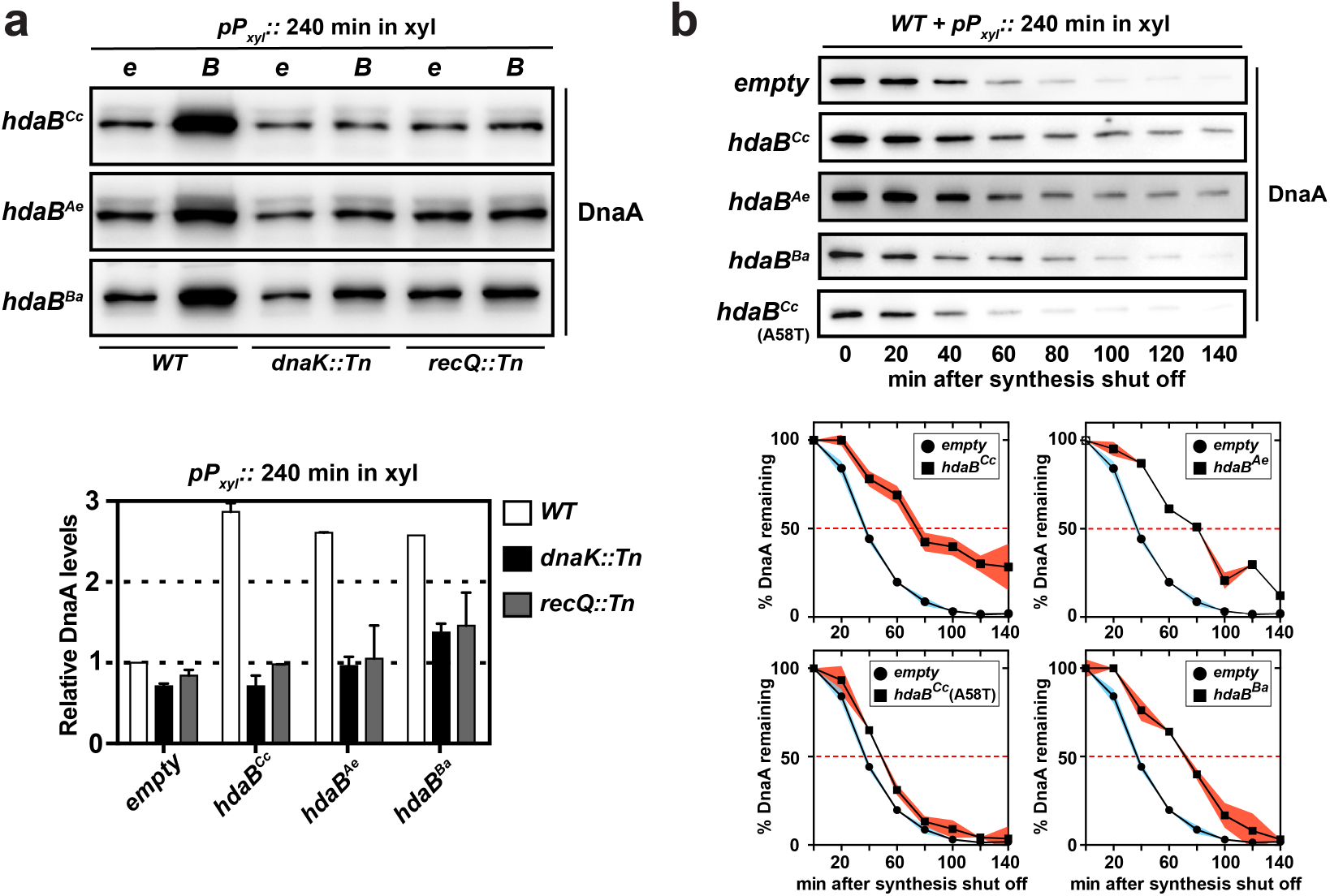
HdaB proteins stabilize DnaA in *C. crescentus*. **(a)** Relative levels of DnaA in *hdaB*-expressing cells. NA1000 (*WT*), JC2080 (*dnaK::Tn*) or JC2079 (*recQ::Tn*) cells carrying the empty (e) control vector (pBXMCS-4) or derivatives (B) expressing *hdaB^Cc^*, *hdaB*^*Ae*^ or *hdaB*^*Ba*^ from the *Pxyl* promoter were cultivated to mid-exponential phase in PYEG medium and xylose was added for the last 4 hours. Cell extracts were then prepared for immunoblotting using anti-DnaA antibodies. Images of representative blots are shown in the upper panel. Relative quantifications of DnaA levels are shown in the lower panel. Error bars correspond to standard deviations between three independent experiments. **(b)** DnaA stability upon expression of *hdaB* homologs. NA1000 (*WT*) cells carrying the empty control vector (pBXMCS-4) or derivatives expressing *hdaB^Cc^*, *hdaB*^*Ae*^, *hdaB*^*Ba*^ or *hdaB*^*Cc*^*(A58T)* from the *Pxyl* promoter were cultivated to mid-exponential phase in PYEG medium and xylose was added for 4 hours before chloramphenicol (2 μg/ml) was added to shut-off protein synthesis. Cell extracts were then prepared at the indicated time points for immunoblotting using anti-DnaA antibodies. Images of representative blots are shown in the upper panel. Relative quantifications of DnaA levels at each time point are shown in the lower panel. Red- or blue-shaded areas correspond to standard deviations between three independent experiments.

Collectively, this data supports a model where HdaB interacts with HdaA to block the RIDA process and stimulate the initiation of DNA replication by DnaA.

### Mutations in *dnaK* and *recQ* can bypass the toxicity of HdaB

As described earlier, most cells expressing HdaB proteins rapidly loose viability when actively growing in liquid or on solid (Fig. 2 and Supplementary Fig. 2) media. If this toxicity is due to the over-initiation of DNA replication (Fig. 3), as anticipated if the target of HdaB is HdaA (Fig. 4), one would expect that suppressors of this toxicity should carry mutations that reduce the levels of active DnaA and/or that reduce initiation frequency and/or that restore viability despite excessive chromosome replication ^21^. To test if this was indeed the case, we isolated random transposon mutants that could bypass the toxicity associated with the expression of HdaB^Ae^ from *Pxyl* on a medium copy number vector in *C. crescentus* (Fig. 2b). Out of a total of ∼8’000 mutants that were screened (Supplementary Fig. 4), two extragenic suppressors acquired the capacity to grow and form colonies despite the presence of the xylose inducer (Fig. 2 and Supplementary Fig. 2).

The first mutant carried a transposon insertion in the *dnaK* gene (Supplementary Fig. 5). Strikingly, *dnaK* mutations had already been randomly isolated previously and shown to bypass the lethality associated with the over-production of DnaA in *C. crescentus* ^12^. DnaK is an essential chaperone activated by its non-essential co-chaperone DnaJ ^22,23^. DnaKJ inhibits the transcription of *lon*, itself encoding the main protease that degrades DnaA during exponential phase ^12^. In the mutant we isolated, the transposon was inserted in the disruptable 3’ region of the *dnaK* open reading frame (ORF) (Supplementary Fig. 5) located upstream of the co-transcribed *dnaJ* ORF ^22,24^. Thus, the transposon most likely decreases DnaK activity and shuts off *dnaJ* transcription through a polar effect. Indeed, the use of a transcriptional reporter to compare the activity of the *lon* promoter showed that it was ∼2-fold more active in the *dnaK::Tn* than in the wild-type strain (Supplementary Fig. 6). Consistent with this finding, immunoblotting experiments then revealed that the *dnaK::Tn* mutation prevented the excessive accumulation of DnaA in *C. crescentus* cells expressing HdaB^Cc^, HdaB^Ae^ or HdaB^Ba^ (Fig. 5), most likely by stimulating DnaA degradation by Lon. Finally, qPCR experiments to measure the *Cori/ter* ratio (Fig. 3c) and microscopy experiments (Fig. 2 and Supplementary Fig. 2) revealed that the isolated *dnaK::Tn* mutation largely attenuated the over-initiation and the cell elongation phenotypes associated with the ectopic expression of HdaB proteins in *C. crescentus*. Altogether, these results are then fully consistent with the proposal that actively growing *C. crescentus* cells die as a consequence of elevated DnaA levels when HdaB proteins are expressed.

The second suppressor mutant carried a transposon insertion in the *recQ* gene encoding a DNA helicase involved in DNA repair ^25^. Interestingly, RIDA deficiencies in *E. coli* were shown to provoke cell death through the accumulation of DNA strand breaks connected with the accumulation of reactive oxygen species (ROS) ^21,26,27^ and RecQ has been shown to be necessary for such ROS-dependent death as also seen during the so-called “thymineless death” (TLD) process ^28,29^. Then, we tested whether *C. crescentus* cells over-expressing HdaB proteins accumulated more double-strand breaks (DSB) than control cells using a RecA-CFP reporter expressed from the endogenous *Pvan* promoter and fluorescence microscopy. As expected if HdaB inhibits the RIDA process, we found that cells over-expressing HdaB^Cc^ proteins displayed RecA-CFP foci far more frequently than control cells (∼36% of the cells instead of ∼2% in Supplementary Fig. 7), indicating that potentially lethal DSBs accumulate. More surprisingly, we found that the *recQ::Tn* mutation when HdaB^Cc^ is expressed in exponential phase lead to an inhibition of DnaA accumulation (Fig. 5a) and to a reduction of the over-initiation (Fig. 3c) and lethal (Fig. 2 and Supplementary Fig. 2b) phenotypes, in a manner very similar to the *dnaK::Tn* mutation. This observation suggested that RecQ and DnaKJ may promote DnaA accumulation through the same Lon-dependent pathway. In agreement with this proposal, *lon* transcription was as much induced in *recQ::Tn* or Δ*recQ* mutants than in the *dnaK::Tn* mutant (Supplementary Fig. 6b). Noteworthy, we also found that RecQ promotes the accumulation of another well-known substrate of the Lon protease: the CcrM DNA methyltransferase (Supplementary Fig. 6c). Thus, this lucky finding also uncovered the existence of a novel RecQ-dependent pathway controlling Lon-dependent degradation in *C. crescentus*.

## DISCUSSION

Lag phase is the most poorly understood phase of the bacterial growth cycle and how bacteria can leave this phase to start proliferating remains obscure. In this study, we identified a novel and conserved small protein that shows similarities with DnaA domain IV (Fig. 1b). We found that HdaB^Cc^ interacts with HdaA^Cc^ (Fig. 4 and Supplementary Fig. 3), most likely titrating HdaA^Cc^ away from DnaA^Cc^ and leading to a transient inhibition of the RIDA process when HdaB^Cc^ is present (Fig. 6). Thus, the ectopic expression of *hdaB* genes in exponentially growing cells promotes DnaA activity (Fig. 3) and represses DnaA degradation by Lon (Fig. 5), leading to over-initiation events and cell death in *C. crescentus* (Fig. 2 and Supplementary Fig. 2). Supporting this model, we found that HdaB toxicity can be bypassed by suppressor mutations in *dnaK* or *recQ* that stimulate Lon-dependent degradation (Fig.5 and Supplementary Fig. 6), restoring normal DnaA levels and eliminating uncontrolled initiation events (Fig. 2, 3).

**Figure 6:**
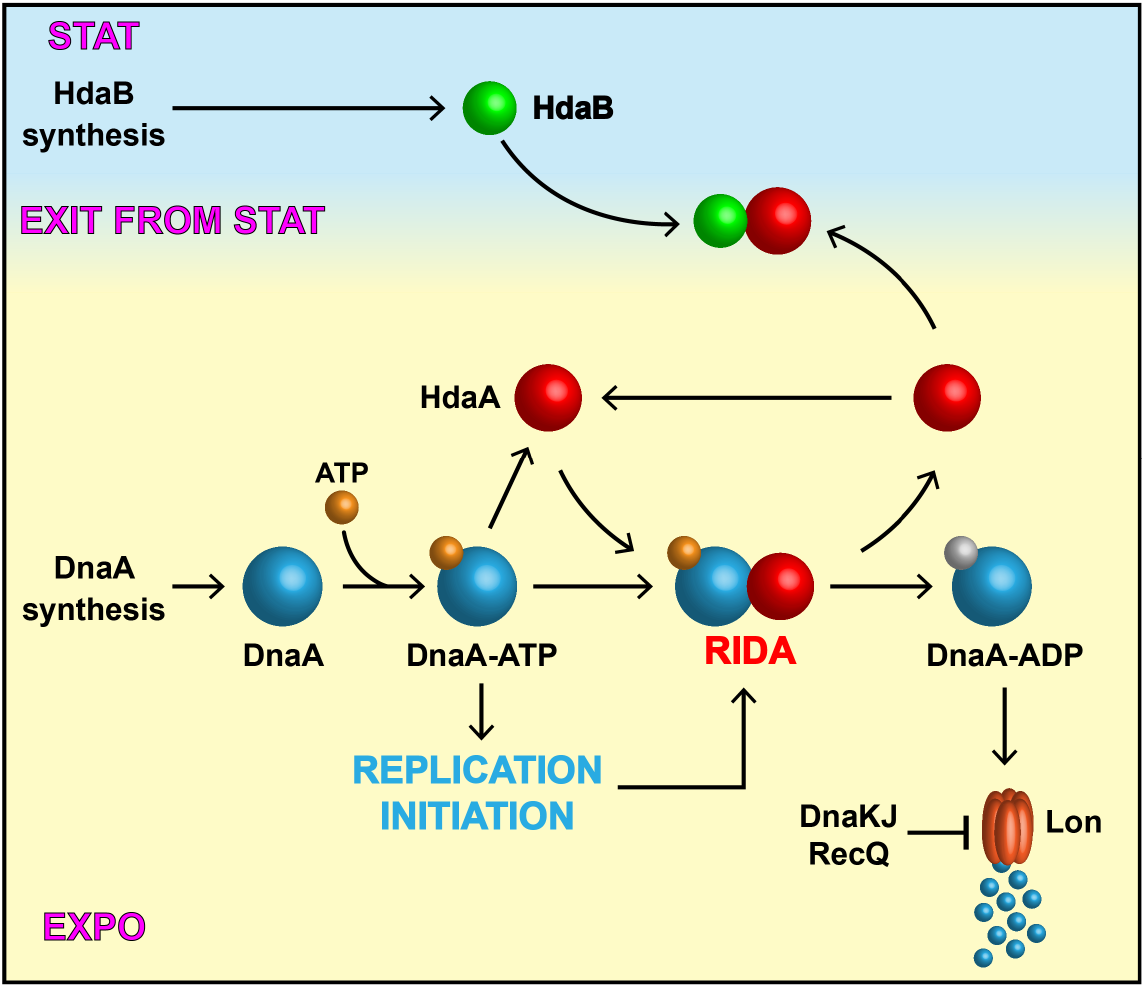
Model for the biological function of HdaB in *C. crescentus.* DnaA synthesis mostly occurs during exponential phase and DnaA associates with ATP soon after its synthesis, giving an active DnaA-ATP complex that can initiate chromosome replication. Once replication has started, HdaA binds to DnaA and stimulates its ATPase activity. The resulting DnaA-ADP complex is actively degraded by the Lon protease as part of the Regulated Inactivation and degradation of DnaA (RIDA) process. Upon entry into stationary phase, DnaA synthesis is strongly repressed, leading to an inhibition of replication initiation and a cell cycle arrest in G1 or G2 phase. Meanwhile, HdaB accumulates through a stimulation of its synthesis. During exit form stationary phase, DnaA synthesis restarts while the RIDA process is inhibited by a sequestration of HdaA by HdaB, leading to optimum levels of DnaA-ATP for rapid replication restart.

### A role for HdaB during exit from stationary phase?

Two observations pointed to a potential role of HdaB proteins during exit from stationary phase (Fig. 6). First, *hdaB*^*Cc*^ transcription (Fig. 1d) and HdaB^Cc^ accumulation (Fig. 1c) are mostly detectable in stationary phase cells. Second, the growth of the *ΔhdaB*^*Cc*^ mutant appears to be delayed during exit from stationary phase, compared to wild-type *C. crescentus* cells (Fig.1e), coincident with a delay in replication restart (Fig.1f). Supporting this last finding, we also observed that the proportion of *cfp-parB* cells displaying two CFP-ParB foci did not increase as much in the absence of *hdaB*^*Cc*^ than in wild-type cells during lag phase (Supplementary Fig. 8), further indicating that HdaB stimulates origin duplication during exit from stationary phase. Altogether, these results suggest that natural amounts of HdaB^Cc^ present in lag phase cells block the RIDA process to promote the activity and the stability of newly synthesized DnaA molecules, stimulating replication restart during exit from stationary phase (Fig. 6).

### Functional conservation for HdaB proteins

HdaB homologs are found in a majority of *Alphaproteobacteria* (Fig. 1a), suggesting that they play an important role in this class of bacteria, which includes a large number of pathogens ^30^. It is possible that HdaB contributes to infectious processes, notably when intracellular pathogens must restart their cell cycle following internalization ^31^. Here, we provide evidence that HdaB homologs from three distantly-related *Alphaproteobacteria* (Fig. 1a), including the *B. abortus* pathogen, can promote DnaA accumulation (Fig. 5) and replication initiation in *C. crescentus* (Fig. 3), indicating a strong functional conservation. Importantly, every bacterium that has a HdaB protein also has a HdaA protein (Fig. 1a), consistent with the proposal that HdaB targets HdaA to stimulate the RIDA process.

### *C. crescentus* death during over-initiation

Similar to what was previously observed when the RIDA process was inhibited by depleting *hdaA*^*Cc*^ ^8^ or by over-expressing a hyper-active DnaA^Cc^(R357A) protein ^6,9^, the ectopic expression of *hdaB* genes in *C. crescentus* lead to over-initiation events coincident with cell division defects and cell death (Fig. 2, 3 and Supplementary Fig. 2). The accumulation of multiple origins perturbed the subcellular localization of ParB (Fig. 3a, b), itself disturbing the bi-polar recruitment of MipZ (data not shown) during S-phase, which plays a critical role in Z-ring positioning and cell division ^19,32^. Cell constriction might be further inhibited by an SOS response activated in response to RecA recruitment to ssDNA. Indeed, we found that many cells displayed RecA-CFP foci (Supplementary Fig. 7) before they died (Fig. 2), indicating the presence of ssDNA zones/lesions on the chromosome prior to cell death ^33^. Such regions can become substrates for the production of double-stranded DNA breaks (DSB) by reactive oxygen species (ROS) leading to cell death in bacteria ^21,26^. Interestingly, such rapid death in response to replication defects is reminiscent of the so-called thymineless death (TLD) process that underlies the action of several bactericidal drugs such as trimethoprim or sulfamethoxazole ^29,34,35^.

### A novel regulatory function for the RecQ helicase

It was recently shown that RecQ is required for TLD in *E. coli* ^28,29^. However, in our study, we found that *recQ* mutations did not rescue over-initiating *C. crescentus* cells by increasing their tolerance for over-initiation, but instead, by reducing replication initiation (Fig. 3c and Fig. 5a). Further characterization of the process showed that RecQ promotes DnaA accumulation not only in *hdaB* over-expressing cells (Fig. 5), but also in wild-type cells cultivated in exponential phase (Supplementary Fig. 6c), through a repression of *lon* transcription (Supplementary Fig. 6b). This newly discovered regulatory process has not only an impact on DnaA levels, but also on the levels of other Lon substrates such as the cell cycle dependent DNA methyltransferase CcrM (Supplementary Fig. 6) which plays a key role in the network controlling the cell cycles of *Alphaproteobacteria ^36,37^*. Understanding how RecQ inhibits Lon-dependent proteolysis and may affect genome maintenance or cell cycle progression in *Alphaproteobacteria* provides a very interesting outlook for this study.

## SUPPLEMENTARY DATA

Supplementary data are available.

## ACKNOWLEDGEMENTS

We thank Laurent Casini for his help during the genetic screen, Regis Hallez for advices to perform immunoprecipitation assays and the Genomic Technology Facility of the University of Lausanne for advices for qPCR analysis. We thank members of the Collier/Viollier/Genevaux/Falquet laboratories for helpful discussions during the project. Finally, we thank Stefano Sanselicio, Clément Gallay, Tiancong Chai and Laurent Casini for their feedback on the manuscript.

## FUNDING

This work was supported by the Swiss National Science Foundation [CRSII3_160703 to JC].

## CONFLICT OF INTEREST

None

