## Supplementary Information for "HdaB: a novel and conserved DnaA-related protein that targets the RIDA process to stimulate replication initiation"

### Content:

Supplementary Figures with legends (page 2)

Supplementary Materials and Methods (page 11)

Supplementary tables describing oligonucleotides, plasmids and strains used in this study (page 16)

Supplementary references (page 19)

### SUPPLEMENTARY FIGURES WITH LEGENDS

### Figure S1

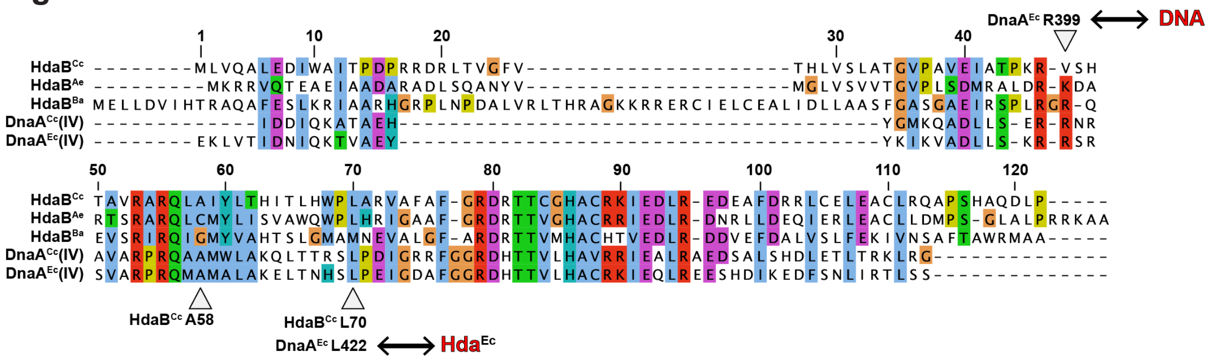


**Figure S1: Similarities between selected HdaB proteins and DnaA domains IV.** Amino-acid sequences of HdaB homologs or DnaA domains IV were downloaded from NCBI for strains NA1000 (*C. crescentus*), CB48 (*A. excentricus*), 2308 (*B. abortus*) or MG1655 (*E. coli*) and alignments were done using the MUSCLE software[1]. For DnaA$^{Cc}$ and DnaA$^{Ec}$, domains IV span I368-G490 and E374-S467, respectively. Residues of particular interest are indicated with arrows.

### Figure S2

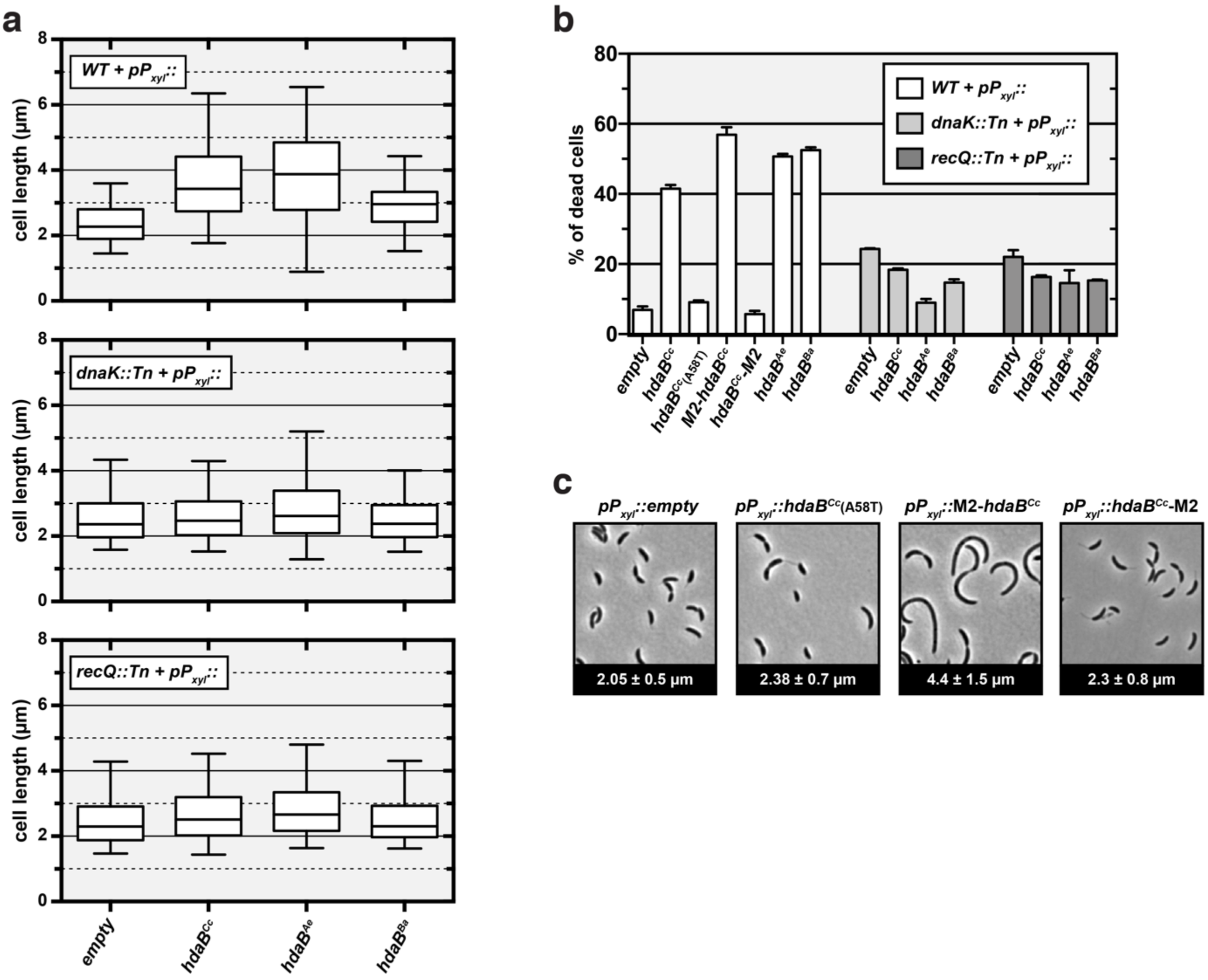


**Figure S2: *hdaB* expression interferes with cell division and leads to cell death. (a)** Cell size distributions upon expression of *hdaB* genes. NA1000 (*WT*) cells carrying pBXMCS-4 (empty) or the indicated pBX-HdaB plasmids were cultivated into PYEG until mid-exponential phase and xylose was then added into the medium for 4 hours before microscopy analysis. Ph3 images were then analyzed with ImageJ to estimate the length of minimum 1000 cells per strain. The line at the center of each box indicates the median medial axis length of cells (in µm). The limits of the box are the first and third quartiles. The whiskers are the 5% and 95% quantiles. **(b)** Proportion of dead cells following expression of *hdaB* genes. NA1000 (WT), JC2080 (*dnaK::Tn*) or JC2079 (*recQ::Tn*) cells carrying pBXMCS-4 (empty) or pBX-HdaB plasmids were cultivated in PYEG to mid-exponential phase and xylose was then added into the medium for 4 hours before LIVE/DEAD staining and fluorescence microscopy imaging. Cells giving a signal for SYTO9 and PI were considered as dead and are reported here as a percentage of the total number of cells (minimum 1000 cells per strain). Error bars correspond to standard deviations from three independent experiments. **(c)** Cellular morphology of *C.*

*crescentus* cells expressing modified *hdaB* genes. NA1000 (*WT*) cells carrying pBXMCS-4 (empty) or pBX-HdaB derivatives were cultivated into PYEG medium until mid-exponential phase and xylose was added for 4 hours before microscopy images were acquired. The numbers indicated at the bottom of each image correspond to the median medial axis length of cells in μm with standard deviations from three independent experiments.

### Figure S3

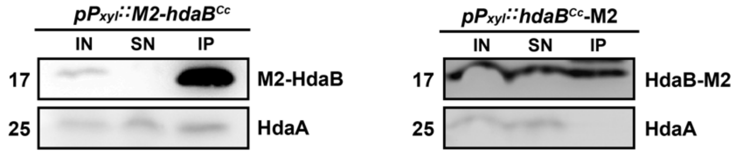


**Figure S3: HdaB interacts with HdaA in *C. crescentus*.** NA1000 (*WT*) cells carrying pBX-M2-HdaB$^{Cc}$ or pBX-HdaB$^{Cc}$-M2 were cultivated to mid-exponential phase in PYEG medium. Xylose was then added into the medium for 4 hours (for *M2-tagged hdaB*$^{Cc}$ expression) before cell lysis and immunoprecipitation using anti-M2 antibodies. Images show immunoblots obtained using anti-HdaA or anti-M2 antibodies. IN: cell extract/input; SN: non-precipitated proteins/supernatant; IP: precipitated proteins/immunoprecipitation. Protein bands shown on these images correspond to proteins at the expected sizes of M2-tagged HdaB$^{Cc}$ (17kD) or HdaA$^{Cc}$ (25kD).

### Figure S4

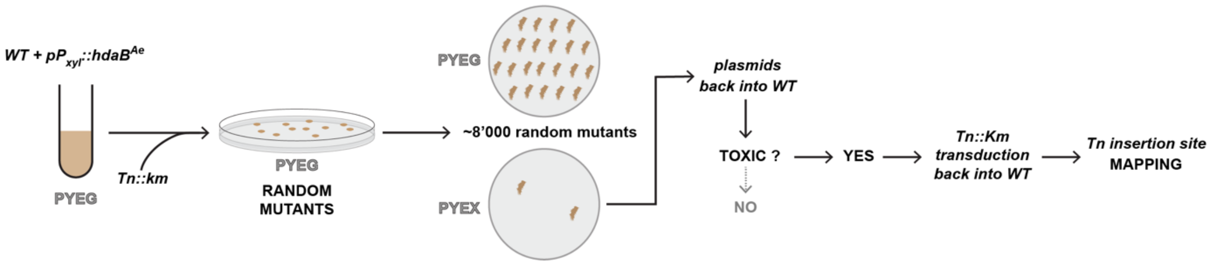


**Figure S4: Schematic describing the genetic screen to isolate *C. crescentus* suppressors that can survive despite *hdaB*$^{Ae}$ over-expression.** *Tn::kan* stands for the *himar1* transposon used for this screen.

### Figure S5

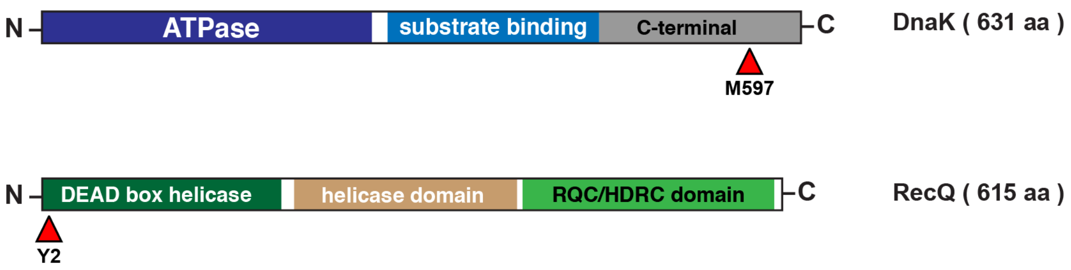


**Figure S5: Schematic showing where transposons were inserted into *dnaK* and *recQ* genes in strains JC2080 and JC2079.** In strain JC2080, the *himar1* transposon was found inserted at position 9955 (into codon encoding M597) of the NA1000 genome available on NCBI. In strain JC2079, the *himar1* transposon was found inserted at position 3735008 (into codon encoding Y2) of the NA1000 genome available on NCBI. aa stands for amino-acids.

### Figure S6

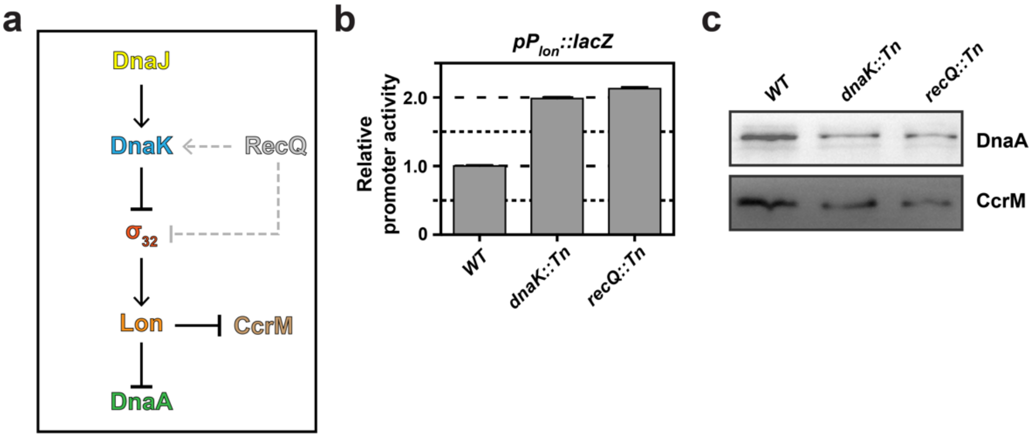


**Figure S6: RecQ and DnaK/J stimulate Lon-dependent degradation in *C. crescentus*. (a)** Schematic depicting the impact of DnaK/DnaJ on DnaA and CcrM degradation by Lon[2] (solid arrows). Results described in this study also uncovered a novel RecQ-dependent pathway promoting DnaA/CcrM degradation by Lon (dashed light-grey arrows). **(b)** RecQ and DnaK/J repress *lon* transcription. The p*lacZ290*-$P_{lon}$ plasmid was introduced into NA1000 (*WT*), JC2080 (*dnaK::Tn*) and JC2079 (*recQ::Tn*) cells. Cells were then cultivated in PYE medium and β-galactosidase assays were performed when cultures reached mid-exponential phase. Activities (in Miller Units) were normalized so that the activity of $P_{lon}$ in *WT* cells equals 1. Error bars correspond to standard deviations from minimum three experiments. **(c)** DnaK/J and RecQ promote the intracellular accumulation of DnaA and CcrM. Strains NA1000 (*WT*), JC2080 (*dnaK::Tn*) and JC2079 (*recQ::Tn*) were cultivated in PYE medium and samples were collected in mid-exponential phase. Proteins were analyzed by SDS-PAGE and detected by immunoblotting using an anti-DnaA or anti-CcrM antibodies.

### Figure S7

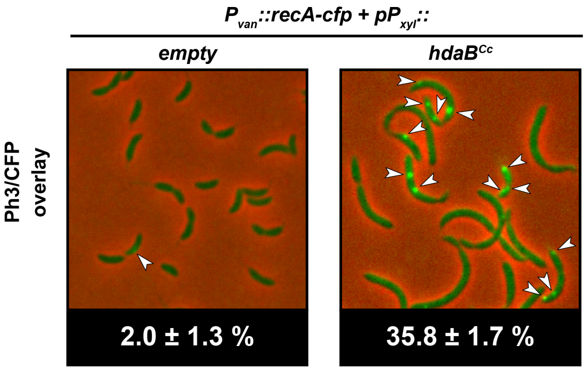


**Figure S7: Expression of *hdaB*$^{Cc}$ promotes RecA-CFP foci formation in *C. crescentus*.** JC2086 (*Pvan::recA-cfp*) cells carrying pBXMCS-4 (empty) or pBX-HdaB$^{Cc}$ were cultivated in PYEG to mid-exponential phase. Xylose and vanillate were then added into the medium for 4 and 5.5 hours, respectively, prior to microscopy imaging. Representative overlays of Ph3 and CFP fluorescence images are shown. The percentages of cells displaying minimum one RecA-CFP focus was estimated using ImageJ on minimum 1000 cells and is indicated under each image (standard deviations from three independent experiments are also included).

**Figure S8**

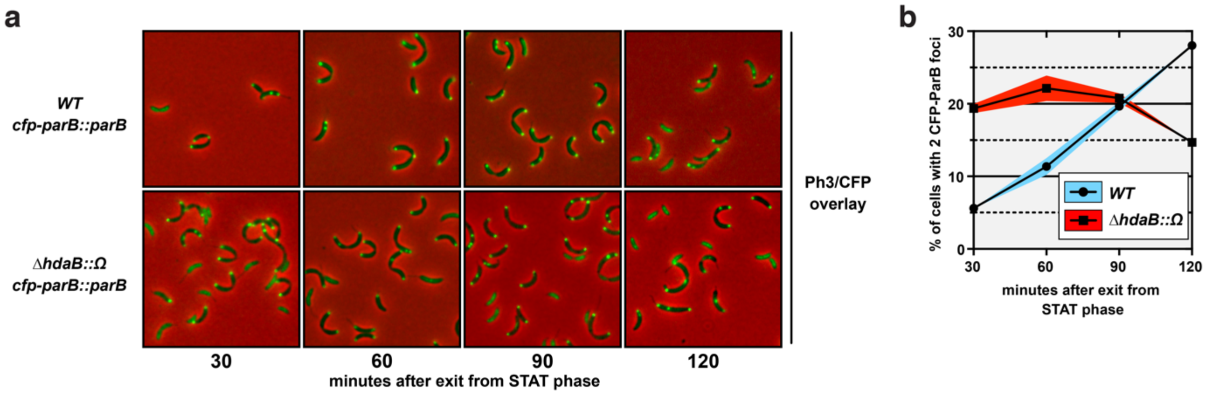


**Figure S8: Replication restart is delayed in Δ*hdaB::Ω* cells.** Strains MT190 (*cfp::parB*) and JC2084 (Δ*hdaB::Ω cfp::parB*) were cultivated in PYE medium until stationary phase (overnight) and cultures were diluted back into fresh PYE medium to reach an $OD_{660nm}$ of 0.1 at time 0. Cells were analyzed by fluorescence microscopy every 30 min. after for 120 min. **(a)** Microscopy images showing Ph3/CFP overlays to visualize CFP-ParB foci in single cells. **(b)** The number of CFP-ParB foci per cell was estimated using ~1000 cells acquired using each strain at the indicated time points. Results are shown as a percentage of cells displaying two CFP-ParB foci in each population. The blue and the red shaded areas correspond to the standard deviations from two independent experiments.

### SUPPLEMENTARY MATERIAL AND METHODS

#### Identification of *hdaB* homologs in *Alphaproteobacteria*

Homologs of *hdaB*$^{Cc}$ were identified using a dataset comprising genomes of one selected species of each genus of *Alphaproteobacteria* as described in [3]. Presence or absence of *hdaB* genes was defined using bidirectional best blast hit (BBH) criterion with an e-value of 0.0001 for the Blast analysis.

#### Plasmid constructions

All PCR amplifications used in cloning procedures were done using *Caulobacter crescentus* NA1000 genomic DNA isolated using the PurGene genomic DNA extraction kit (QIAGEN, DE) and the KOD hot start DNA polymerase (Novagen, DE). DNA sequences of plasmid inserts were systematically confirmed by DNA sequencing. *Escherichia coli* Top10 was used for standard cloning procedures.

*Construction of pBX-HdaB derivatives*

The coding sequence of *hdaB*$^{Cc}$ (genome NC_011916; nucleotides: 1217028-1217396) was amplified using primers AF11 and AF12, digested with NdeI/EcoRI and ligated into NdeI/EcoRI-digested pBXMCS-4 to construct **pBX-HdaB$^{Cc}$**. **pBX-HdaB$^{Cc}$(A58T)** was accidently obtained during the cloning procedure. Synthetic DNA fragments (Integrated DNA Technologies-IDT, Coralville, IA, USA) described in Supplementary note were amplified using M13_FWD and Van [4] primers, digested with NdeI/EcoRI and ligated into NdeI/EcoRI-digested pBXMCS-4 to construct **pBX-M2-HdaB$^{Cc}$**, **pBX-HdaB$^{Cc}$-M2, pBX-HdaB$^{Ae}$ and pBX-HdaB$^{Ba}$**.

*Construction of pBV-M2-HdaB$^{Cc}$ and pBV-HdaB$^{Cc}$-M2*

The M2-HdaB$^{Cc}$ and HdaB$^{Cc}$-M2 inserts from pBX-M2-HdaB$^{Cc}$ and pBX-HdaB$^{Cc}$-M2 were excised by restriction using NdeI/EcoRI and cloned into NdeI/EcoRI-digested pBVMCS-4 to construct **pBV-M2-HdaB$^{Cc}$** and **pBV-HdaB$^{Cc}$-M2**, respectively.

*Construction of pV-RecA-CFP*

The coding sequence of *recA* (genome NC_011916; nucleotides: 1245163-1246233) was amplified using primers AF87 and AF88, digested with NdeI/EcoRI and ligated into NdeI/EcoRI-digested pVCFPC-1 to construct **pV-RecA-CFP**.

*Construction of placZ290 derivatives*

The promoter regions of *hdaB*$^{Cc}$ (genome NC_011916; nucleotides: 1216700-1217153 = 454 bp) and *lon* (genome NC_011916; nucleotides: 2183418-2183911 = 493 bp) were amplified using primers pairs AF18/AF19 or AF101/102, digested with EcoRI/XbaI and ligated into I-EcoRI/XbaI-digested *placZ290* to construct ***placZ290*-P$_{hdaB}$** and ***placZ290*-P$_{lon}$**, respectively.

*Construction of pNPTS138-ΔhdaB*

The 550/600 pb sequences upstream and downstream of the *hdaB*$^{Cc}$ coding sequence were amplified using primer pairs AF20/AF21 or AF22/23, and digested by HindIII/BamHI or BamHI/EcoRI, respectively. Both fragments were then ligated into HindIII/EcoRI-digested pNPTS138 to construct **pNPTS138-Δ*hdaB***.

*Construction of pNPTS138-ΔhdaB' and pNPTS138-ΔhdaB::Ω*

The 519/524 bp sequences upstream and downstream of the *hdaB*$^{Cc}$ coding sequence were amplified using primer pairs JC41/JC42 or JC43/JC44, and digested by HindIII/BamHI or BamHI/NheI, respectively. Both fragments were then ligated into HindIII/NheI-digested pNPTS138 to construct **pNPTS138-Δ*hdaB'***. The Ω cassette was excised from pBOR using BamHI digestion and cloned into BamHI-digested pNPTS138-Δ*hdaB'* to construct **pNPTS138-Δ*hdaB::* Ω**

*Construction of pNPTS138-P$_{hdaB}$-M2-hdaB*

A DNA fragment including the *hdaB*$^{Cc}$ promoter region (563 bp) upstream of a sequence (546 bp) encoding an M2-tagged HdaB$^{Cc}$ protein was generated using overlap extension PCR using primers AF71/AF73/AF72/AF74. This fragment was digested by HindIII/EcoRI and cloned into HindIII/EcoRI-digested pNPTS138 to construct **pNPTS138-*P$_{hdaB}$-M2-hdaB***.

**Strain construction**

When necessary, replicating or integrative plasmids were introduced into *C. crescentus* cells by transformation or conjugation, respectively, using standard procedures. *E. coli* S17-1 and S17-1 *λpir* were used for conjugation and transposon delivery/insertion site mapping, respectively.

*Construction of* ***JC1456, JC166*** *and* ***JC2106****:*

To construct strains **JC1456** and **JC2106**, the pNPTS138-Δ*hdaB* or the pNPTS138-$P_{hdaB}$-*M2-hdaB* plasmids were introduced into the NA1000 strain. Plasmids integrated at the *hdaB* region of the chromosome by homologous recombination. The resulting strains were then cultivated in PYE medium until stationary phase before plating onto PYEA supplemented with 3% of sucrose. Single colonies were then transferred in parallel onto PYEA + kanamycin (Km: 25 µg/mL) and PYEA to select $Km^S$ colonies (plasmid re-excision). We then screened for colonies carrying Δ*hdaB* or the $P_{hdaB}$::*M2-hdaB* allele by colony PCR using primers AF50/51 or AF50/91, respectively.

To construct strain **JC166**, the pNPTS138-Δ*hdaB::Ω* plasmid was introduced into the NA1000 strain. The same procedure as described above was then used, except that Δ*hdaB::Ω* colonies were selected directly on spectinomycin/streptomycin-containing plates before genotype confirmation by colony-PCR using primers AF50/51.

*Construction of* ***JC2084****:*

The Δ*hdaB::Ω* allele was transduced into the MT190 strain using ΦCr30-mediated generalized transduction and selecting for spectinomycin/streptomycin-resistant colonies. The genotype was then confirmed by colony-PCR using primers AF50/51.

*Construction of* ***JC2086****:*

Plasmid pV-RecA-CFP was integrated into the *vanA* promoter of strain NA1000 by a single integration event. Integration was verified by colony PCR using primers RecUni-1/RecVan-2 [4].

**Estimation of the proportion of dead cells in populations**

To estimate the proportion of dead cells in *C. crescentus* populations, cultures were centrifuged and cells were resuspended into a mixture of Syto9 and PI (Propidium Iodide) from the LIVE/DEAD *Bac*Light bacterial viability kit (Thermo-Fisher, MA, USA). After 10 minutes of incubation, cells were imaged by fluorescence microscopy. Cells giving a red fluorescent signal (PI staining) were considered as dead.

**SUPPLEMENTARY NOTE**

5’-3’ sequences of synthetic DNA fragments used in this study. Sequences added for amplification using M13_FWD and Pvan-for primers [4] are shown in green colour. Sequences corresponding to restriction sites are shown in bold. Sequences encoding the M2 tag are underlined.

Used for **pBX-M2-HdaB**$^{\mathbf{Cc}}$ (Van_NdeI_$^{\mathit{FLAG-}}$***hdaB***_EcoRI_M13for):

*GACGTCCGTTTGATTACGATCAAGATTGG***CATATG**GATTACAAGGACGATGATGATAAGatgctggttcaagcccttgaggacatctgggcgatcacgcccgatccccgacgcgaccgcctgacggtcggcttcgtcacccaccttgtctccctggccaccggcgtgccggcggtggagatcgcgacgcccaagcgcgtgtctcatacggcggtgcgcgctcggcagctcgccatctacctgacccacatcaccctgcactggcccctggcgcgcgtggccttcgcttttggccgcgatcgcaccacctgcgggcacgcctgccgcaagatcgaggatctgcgcgaggacgaggcctttgaccgccggctctgcgaactggaggcctgcctgcgtcaggccccgtcgcacgctcaggatctgccgtaa**GAATTC***TCGTGACTGGGAAAACCCTGGC*

= *hdaB*$^{Cc}$ coding sequence.

Used for **pBX-HdaB**$^{\mathbf{Cc}}$**-M2** (Van_NdeI_***hdaB***$^{\mathit{-FLAG}}$_EcoRI_M13for):

*GACGTCCGTTTGATTACGATCAAGATTGG***CATATG**ctggttcaagcccttgaggacatctgggcgatcacgcccgatccccgacgcgaccgcctgacggtcggcttcgtcacccaccttgtctccctggccaccggcgtgccggcggtggagatcgcgacgcccaagcgcgtgtctcatacggcggtgcgcgctcggcagctcgccatctacctgacccacatcaccctgcactggcccctggcgcgcgtggccttcgcttttggccgcgatcgcaccacctgcgggcacgcctgccgcaagatcgaggatctgcgcgaggacgaggcctttgaccgccggctctgcgaactggaggcctgcctgcgtcaggccccgtcgcacgctcaggatctgccgGATTACAAGGACGATGATGATAAGtaa**GAATTC***TCGTGACTGGGAAAACCCTGGC*

Used for **pBX-HdaB**$^{\mathbf{Ae}}$ (Van_NdeI_***hdaB***$^{\mathit{Aex}}$_EcoRI_M13for*):*

*GACGTCCGTTTGATTACGATCAAGATTGG***CATATG**aagcgtcgggtccaaacggaggccgagatcgcggcggatgcgcgggcggatctcagccaagccaattatgtgatggggctcgtgagcgtggtgaccggcgtgccgctctccgatatgcgcgcgctggatcggaaggatgcgcggacgtcccgggcccggcaactctgcatgtatctcatctcggtcgcgtggcagtggccgctgcaccggatcggggccgccttcggccgtgatcgtaccacggtggggcacgcctgtcgtcggatcgaagacctgcgtgacaaccgcctgctggatgagcagatcgagcgcctcgaagcgtgcctgctggatatgccctccggcctggcgctgccgcgtcggaaggcggcgt**GAATTC***TCGTGACTGGGAAAACCCTGGC*

= *hdaB*$^{Ae}$ coding sequence (Genome NC_014816; nucleotides 658687-659064).

Used for **pBX-HdaB**$^{\mathbf{Ba}}$ (Van_NdeI_***hdaB***$^{\mathit{Bab}}$_EcoRI_M13for*):*

*GACGTCCGTTTGATTACGATCAAGATTGG***CATATG**gagctgctggacgtcatccacacgcgggcgcaa
gcgtttgagtcgctcaagcgtatcgcggcccgtcacgggcggcccctgaatcccgacgcgctggtccgtctgacgcaccgtgcggg
gaagaagcgtcgtgaacggtgcatcgaactctgcgaagcgctgatcgacctgctggccgcgagcttcggcgcctcgggggcggag
atccgctcgcccctgcgggggcggcaggaggtctcgcgcatccgtcaaatcgggatgtacgtcgcgcatacgtccctcggcatggc
catgaatgaagtggccctggggtttgcccgggaccggaccaccgtgatgcatgcgtgccacaccgtggaggatctccgtgacgatgt
cgagtttgacgccctggtgtccctgttcgagaagatcgtcaactcggcctttaccgcgtggcgtatggcggcgt**gaATTC***TCGT*
*GACTGGGAAAACCCTGGC*

= *hdaB*$^{Ba}$ coding sequence (Genome NC_007618; nucleotides 1536267-1536722).

### SUPPLEMENTARY TABLES

**Supplementary Table S1: Oligonucleotides used in this study.** Restriction enzyme recognition sites used for cloning purposes are in bold.

| AF number | Sequence (5'-3') | Used for |
|---|---|---|
| AF11 | **CATATG**CTGGTTCAAGCCCTTG | pBX-HdaB$^{Cc}$ construction |
| AF12 | **GAATTC**TTACGGCAGATCCTGAGCG | pBX-HdaB$^{Cc}$ construction |
| AF18 | **GAATTC**AGCCCCCTTGTGGCCGT | *placZ290*-$P_{hdaB}$ construction |
| AF19 | **TCTAGA**GCGATCTCCACCGCCGGC | *placZ290*-$P_{hdaB}$ construction |
| AF20 | **AAGCTT**GTGAAGGCCGTGGACAC | pNPTS138-Δ*hdaB* construction |
| AF21 | **GGATCC**CAGCATGGCGTCCTC | pNPTS138-Δ*hdaB* construction |
| AF22 | **GGATCC**GCATGAGGCGACGTTCAG | pNPTS138-Δ*hdaB* construction |
| AF23 | **GAATTC**CGACTTCCCCGCCTGGTTC | pNPTS138-Δ*hdaB* construction |
| AF50 | CTGGGTGATGGCGATACG | Test for deletion of the $hdaB^{Cc}$ |
| AF51 | TCATGCCATCCGGTAGTGTC | Test for deletion of the $hdaB^{Cc}$ |
| AF71 | **AAGCTT**CCGTCAATGTTCACACACCG | pNPTS138-$P_{hdaB}$-*M2-hdaB* construction |
| AF72 | ATGGATTACAAGGACGATGATGATAAGATGCTGGTT CAAGCCCTT | pNPTS138-$P_{hdaB}$-*M2-hdaB* construction |
| AF73 | CTTATCATCATCGTCCTTGTAATCCATGGCGTCCTCT CCTCC | pNPTS138-$P_{hdaB}$-*M2-hdaB* construction |
| AF74 | **GAATTC**TGGCCTCGTCCAGGGTC | pNPTS138-$P_{hdaB}$-*M2-hdaB* construction |
| AF77 | CGCGGAACGACCCACAAACT | qPCR |
| AF78 | CAGCCGACCGACCAGAGCA | qPCR |
| AF79 | CCGTACGCGACAGGGTGAAATAG | qPCR |
| AF80 | GACGCGGCGGGCAACAT | qPCR |
| AF87 | **CATATG**ACAAGTCAGGCGGC | pV-RecA-CFP construction |
| AF88 | **GAATTC**GTCCTCTTCGCCCTCTTCCG | pV-RecA-CFP construction |
| AF91 | CTTATCATCATCGTCCTTGTAATC | Test for recombinant of integration of pNPTS138-$P_{hdaB}$-*M2-hdaB* |
| AF101 | **GAATTC**GCTGCTGATCTATGCCG | *placZ290*-$P_{lon}$ construction |
| AF102 | **TCTAGA**GAACACAACGATATCCCGC | *placZ290*-$P_{lon}$ construction |
| JC41 | GGT**AAGCTT**TGGACACGACCGGCGCGGGC | pNPTS138-Δ*hdaB'* construction |
| JC42 | CC**GGATCC**CTCAAGGGCTTGAACCAGCATG | pNPTS138-Δ*hdaB'* construction |
| JC43 | CC**GGATCC**GCTCAGGATCTGCCGTAATGAC | pNPTS138-Δ*hdaB'* construction |
| JC44 | CCGG**GCTAGC**CGTCGACGCGGGGCGTGGC | pNPTS138-Δ*hdaB'* construction |
| himup2 | GATATTGCTGAAGAGCTTGGCGGCGAA | *Tn* insertion site mapping |
| RecUni-1 | ATGCCGTTTGTGATGGCTTCCATGTCG | Test of recombinants for integration of a plasmid into the *vanA* or *xylX* locus |
| RecVan-2 | CAGCCTTGGCCACGGTTTCGGTACC | Test for integration into the *vanA* locus |
| Pvan-for | GACGTCCGTTTGATTACGATCAAGATTGG | Amplification of synthetic DNA fragments |
| M13-for | GCCAGGGTTTTCCCAGTCACGA | Amplification of synthetic DNA fragments |

### Supplementary Table S2: Plasmids used in this study.

| **Plasmid Name** | **Description** | **Source** |
|---|---|---|
| pGEM-T Easy | TA-cloning vector | Promega (USA) |
| pHPV414 | Himar1 transposon delivery vector (*oriR6K*) | [5] |
| pBXMCS-4 | Medium copy number vector for xylose-inducible gene expression | [4] |
| pBX-HdaB$^{Cc}$ | *hdaB*$^{Cc}$ under the control of $P_{xyl}$ in pBXMCS-4 | This study |
| pBX-HdaB$^{Cc}$(A58T) | *hdaB*$^{Cc}$*(A58T)* under the control of $P_{xyl}$ in pBXMCS-4 | This study |
| pBX-M2-HdaB$^{Cc}$ | *M2-hdaB*$^{Cc}$ under the control of $P_{xyl}$ in pBXMCS-4 | This study |
| pBX-HdaB$^{Cc}$-M2 | *hdaB*$^{Cc}$*-M2* under the control of $P_{xyl}$ in pBXMCS-4 | This study |
| pBX-HdaB$^{Ae}$ | *hdaB*$^{Ae}$ under the control of $P_{xyl}$ in pBXMCS-4 | This study |
| pBX-HdaB$^{Ba}$ | *hdaB*$^{Ba}$ under the control of $P_{xyl}$ in pBXMCS-4 | This study |
| pBVMCS-4 | Medium copy number vector for vanillate-inducible gene expression | [4] |
| pBV-M2-HdaB$^{Cc}$ | *M2-hdaB*$^{Cc}$ under the control of $P_{van}$ in pBVMCS-4 | This study |
| pBV-HdaB$^{Cc}$-M2 | *hdaB*$^{Cc}$*-M2* under the control of $P_{van}$ in pBVMCS-4 | This study |
| pVCFPC-1 | Vector for generating C-terminal protein fusions encoded at the *vanA* locus | [4] |
| pV-RecA-CFP | *recA-cfp* under the control of $P_{van}$ in pVCFPC-1 | This study |
| *placZ290* | Low copy number plasmid to create transcriptional fusions with *lacZ* | [6] |
| *placZ290*-P$_{hdaB}$ | *lacZ* gene under the control of $P_{hdaB}$ in p*lacZ290* | This study |
| *placZ290*-P$_{lon}$ | *lacZ* under the control of $P_{lon}$ in p*lacZ290* | This study |
| pNPTS138 | Suicide vector carrying the *sacB* gene | D. Alley, unpublished |
| pNPTS138-Δ*hdaB* | pNPTS138 derivative used to create the □*hdaB* in-frame deletion strain | This study |
| pNPTS138-$P_{hdaB}$-*M2-hdaB* | pNPTS138 derivative used to create replace *hdaB* by *M2-hdaB* | This study |
| pBOR | Vector carrying a Spectinomycin/Streptomycin cassette (Ω derivative of pHP45Ω) | [7] |
| pNPTS138-Δ*hdaB'* | pNPTS138 derivative used to construct pNPTS138-Δ*hdaB::* Ω | This study |
| pNPTS138-Δ*hdaB::* Ω | pNPTS138-□*hdaB* derivative used to create the Δ*hdaB::* Ω strain | This study |

### Supplementary Table S3: Bacterial strains used in this study.

| Strain | Genotype | Source |
|---|---|---|
| ***Escherichia coli*** | | |
| Top10 | *F- mcrA Δ(mrr-hsdRMS-mcrBC) φ80lacZΔM15 ΔlacX74 nupG recA1 araD139 Δ(ara-leu)7697 galE15 galK16 rpsL(Str$^R$) endA1 λ$^-$* | Invitrogen (USA) |
| S17-1 *λpir* | *RP4-2(Km::Tn7,Tc::Mu-1) pro-82 λpir recA1 endA1 thiE1 hsdR17 creC510* | [8] |
| S17-1 | *294::RP4-2(Tc::Mu)(Km::Tn7)* | [8] |
| ***Caulobacter crescentus*** | | |
| NA1000 | Synchronizable derivative of wild-type strain CB15 (CB15N) | [9] |
| JC1456 | NA1000 Δ*hdaB* | This study |
| JC166 | NA1000 Δ*hdaB::Ω* | This study |
| JC2106 | NA1000 $P_{hdaB}$ *::M2-hdaB* | This study |
| JC2077 | NA1000 *dnaK::Tn* pBX-HdaB$^{Ae}$ (from genetic screen) | This study |
| JC2078 | NA1000 *recQ::Tn* pBX-HdaB$^{Ae}$ (from genetic screen) | This study |
| JC2080 | NA1000 *dnaK::Tn* | This study |
| JC2079 | NA1000 *recQ::Tn* | This study |
| MT190 | NA1000 *cfp-parB::parB* | [10] |
| JC208 | NA1000 $P_{xylX}$*::hdaA-gfp* | [11] |
| JC2086 | NA1000 $P_{vanA}$*::recA-cfp* | This study |
| JC2084 | MT190 Δ*hdaB::Ω* | This study |
